# Structural connectome architecture and biological vulnerability shape cortical atrophy in cocaine use disorder

**DOI:** 10.1101/2025.08.11.669789

**Authors:** Ziteng Han, Tiantian Liu, Kexin Wang, Guoyuan Yang, Tianyi Yan

## Abstract

Cocaine use disorder (CUD) is prevalent and characterized by widespread gray matter atrophy across the cerebral cortex. However, it remains unclear whether and how connectome-based circuits and biological features shape these structural abnormalities. Here, we show that CUD-associated regional atrophy is constrained by the white matter (WM) structural connectome. Along these WM pathways, regions that share similar haemodynamic activity and molecular profiles are more likely to exhibit analogous atrophy patterns. By integrating the structural connectome with these multiple connectivity blueprints, we subsequently identify CUD epicenters and reveal that the prefrontal and visual cortices serve as core systems. Furthermore, we link these epicenters to cortical gene expression patterns and delineate CUD-related transcriptomic enrichment in biological processes, including inter-neuronal communication, synaptic function and neural homeostasis. Finally, we demonstrate that the spatial distribution of these epicenters correlates with cocaine craving-response maps derived from repeated transcranial magnetic stimulation and can track individual variations in clinical behavioural representations, suggesting their potential as key targets for therapeutic intervention. Together, our findings establish a structurally constrained framework for the spread of pathology underlying cortical atrophy in CUD, where initial perturbations propagate via structural connectome pathways to vulnerable regions shaped by neural activity and molecular landscapes.

## Introduction

Cocaine use disorder (CUD), characterized by compulsive cocaine consumption, is considered a global health problem that contributes to significant socioeconomic burdens^1,2^. In particular, CUD is accompanied by psychiatric consequences, with patients exhibiting behavioural alterations in reward processing and impulsivity, as well as experiencing depressive episodes and even suicidal ideation^3–6^. Currently, standard pharmacological and psychosocial treatments are limited by high relapse rates, thus highlighting the need for more effective interventions^7^. Identifying robust and clinically useful neuroimaging-based biomarkers holds promise for advancing our understanding of the pathophysiology of CUD and facilitating the development of reliable therapeutic strategies^8,9^.

Prior neuroimaging studies have investigated brain structural alterations in CUD at the group-level, indicating that cocaine dependence is associated with lower gray matter volume^10,11^. Compared with healthy controls, patients with CUD also exhibit cortical thinning, especially in regions involved in the executive regulation of reward and attention^12^. Extending these findings, a recent mega-analysis revealed that some cortical thinning effects were common across multiple substances (e.g., in the insular cortex), whereas others appeared to be specific to cocaine use (e.g., in the right supramarginal gyrus)^13^. However, although widespread cortical abnormalities have been observed in CUD, the existing results are heterogeneous and their network-level mechanisms remain largely unclear. Understanding the network underpinnings of morphological abnormalities may provide insights into the brain-wide neural substrates of CUD and facilitate effective therapy.

The human brain is a complex network, and inter-regional interactions through axonal pathways scaffold its development and normal function^14,15^. When networked elements are perturbed by pathological processes, the effects propagate among the nodes and infiltrate distributed systems in the brain^16^. Indeed, trans-neuronal spreading of misfolded proteins or inflammatory markers has been documented in multiple neurodegenerative diseases^17–22^. Psychiatric studies also support for the connectome perspective, suggesting that the spatial patterning of neuromorphic deviations is circumscribed by the white matter (WM) architecture^23,24^. Substance dependent-related studies have further revealed evidence that is consistent with this idea. For example, adults with heavy alcohol dependence exhibit lower segregation and greater integration in cortical thickness covariance networks, indicating disruptions in cortico-cortical growth^25,26^. Notably, the spread of pathology can be driven by both WM structural network architecture and local biological vulnerability. Local patterns of hemodynamic activity^27^, transcriptional signatures^28^, neurotransmitter receptor profiles^29^, and cellular composition^30,31^ may influence the vulnerability of neuronal populations to disease. Molecular similarity across regions may serve as a factor promoting disease propagation^32^. Consequently, vulnerable regions are more likely to emerge as disease-related epicenters—that is, regions where the disease process originates or is most prominent—guiding the network spread of pathology and offering insights into potential treatments. Together, these studies raise the possibility that the structural connectome and local biological attributes of the brain jointly shape the spatial pattern of cortical abnormalities in CUD.

In this study, we integrated the brain’s structural connectome with multiple connectivity blueprints to model the network-level pathological process underlying cortical atrophy and investigate the regional heterogeneity of network-level constraints on cortical abnormalities in CUD. More specifically, we tested three inter-related hypotheses. First, we hypothesized that the structural connectome constrains the spread of cortical atrophy across regions in CUD, and the spread of pathology across regions is influenced by their biological feature similarities. Second, we expected that the spread would be amplified in a set of vulnerable brain regions (i.e., disease epicenters) whose connectivity profiles are central to the manifestation of CUD. Third, we hypothesized that the spatial distribution of epicenters is associated with treatment outcomes and can track clinical representation variability.

## Results

### Normative model identifies cortical atrophy in CUD

After quality control, 53 patients (age range:18-48 years, 8 females) with CUD in the discovery cohort, 74 patients (age range:18-50 years, 9 females) with CUD in the replication cohort and 364 healthy controls (age range:18-48 years, 230 females) were included in the present study. We identified cortical thickness deviation patterns for each patient in both the discovery and replication cohorts using a normative modelling approach (**Fig. 1A**). The normative model was trained using multivariate linear regression on cortical thickness data from 364 healthy controls, with age and sex included as covariates. The individualized *W*-score metrics were then averaged within the two independent datasets to generate group-level *W*-score maps. These *W*-scores reflect deviations from normative age- and sex-adjusted predictions, with more negative values corresponding to greater cortical thickness reductions, and more positive values indicating greater cortical thickening. We observed widespread cortical atrophy in patients with CUD, with the most pronounced effects localized in the frontal and visual cortical areas (**Fig. 1B**), which are commonly implicated in cognitive control and sensory processing^33^. In addition, the *W*-score maps derived from the two datasets exhibited significant spatial correlations (**Fig. 1C**; r = 0.38, *P*_spin_ < 0.001), suggesting robust reproducibility across cohorts.

**Fig. 1.**
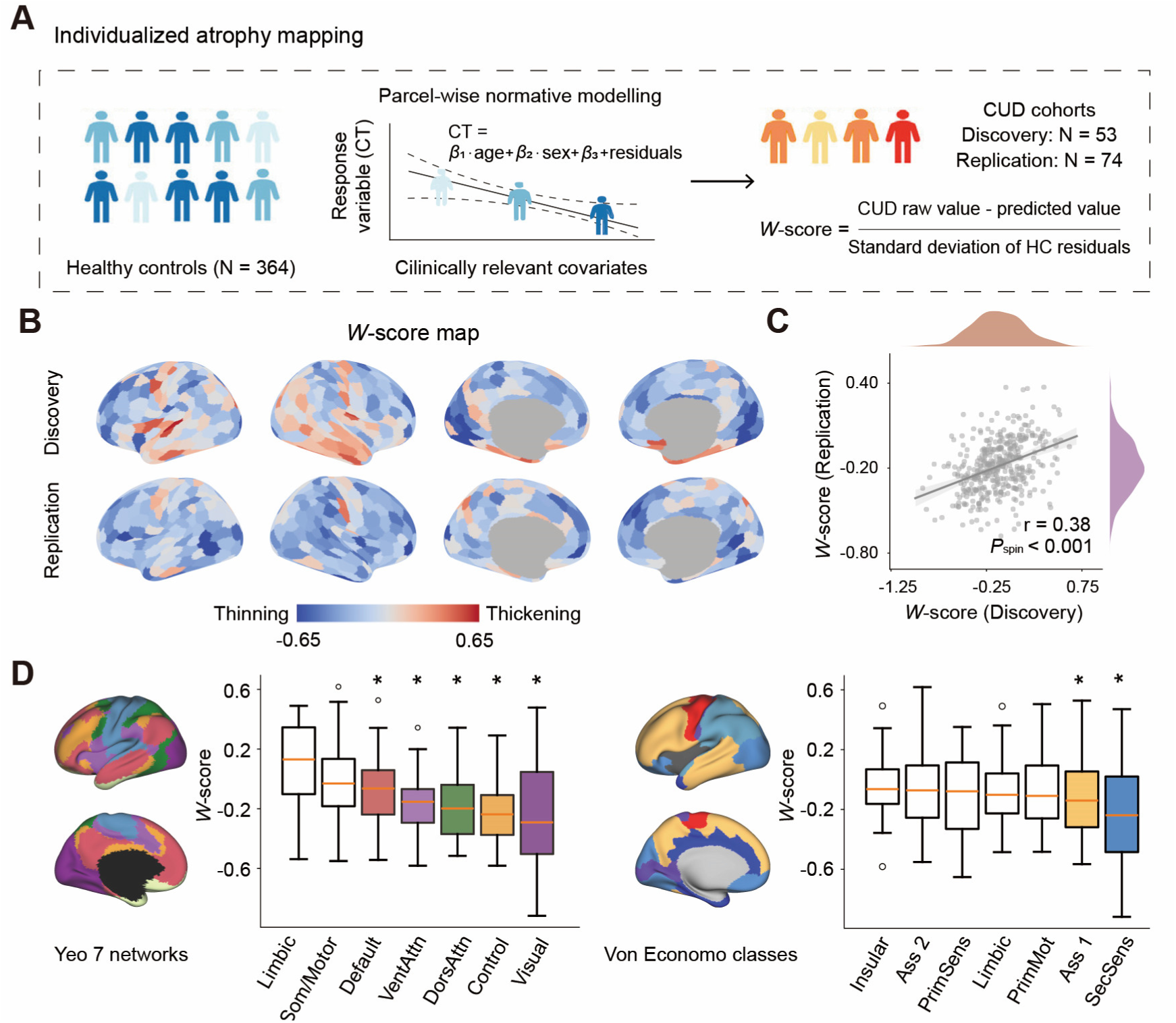
CUD-related cortical thickness deviation patterns derived from normative modelling. (A) We first used a multivariate linear model to fit a normative model for cortical thickness in healthy control participants, controlling for age and sex. The normative model was subsequently applied to the cocaine use disorder (CUD) cohorts to generate individual-level, parcel-wise *W*-score maps, which estimate the cortical thickness deviation from the reference range observed in healthy controls for each patient. (B) Group-level cortical thickness deviation patterns (i.e., *W*-score map) revealed widespread cortical atrophy of CUD patients. (C) The *W*-score maps were reproducible across the two independent cohorts (r = 0.38, *P*_spin_ < 0.001, one-sided). (D) The *W*-score maps in the discovery cohort were stratified into the intrinsic functional networks defined by Yeo et al.^34^ and the cytoarchitectonic classes defined by Von Economo et al.^35^. Boxes represent the interquartile range (IQR), with the median shown as the red horizontal line, while the lower and upper boundaries of each box correspond to the 25th and 75th percentiles, respectively. Asterisks denote statistically significant levels with FDR-corrected *P* < 0.001 (two-sided *t*-test). CT, cortical thickness; Som/Motor, somatomotor network; VentAttn, ventral attention network; DorsAttn, dorsal attention network; Ass, association; PrimSens, primary sensory; PrimMot, primary motor; SecSens, secondary sensory.

To further assess whether these abnormal patterns converge within specific systems, we parcellated each *W*-score map into seven intrinsic functional networks^34^ and canonical cytoarchitectural classes^35^, respectively. We found that atrophy was significantly more enriched in the default mode, frontoparietal control, attention and visual networks, whereas the somatomotor and limbic networks were relatively preserved (**Fig. 1D**; *P*_FDR_ < 0.001). For the Von Economo classes, CUD had greater cortical thinning in the association 1 and secondary sensory classes (**Fig. 1D**; *P*_FDR_ < 0.001). Together, these results demonstrate the sensitivity of the normative modelling approach in detecting atypical cortical changes in CUD, suggesting that CUD-related morphometric alterations are primarily located in the higher-order association systems involved in control, emotional, and decision-making, and the visual systems.

### Structural connectome and local biological vulnerability shape cortical atrophy patterns

Having identified CUD-related cortical atrophy patterns, we next examined whether the spatial distribution of atrophy reflects the structural connectome architecture of the brain. We computed the Pearson correlation coefficient between each brain region’s *W*-score and the average *W*-score of its structurally connected neighbour regions, as defined by a binary normative structural connectome (**Fig. 2A**). A significant positive correlation was observed (**Fig. 2B**; r = 0.42, *P* < 0.001), indicating that regions with great atrophy tend to be structurally connected to other highly atrophied regions. We next assessed the significance of this spatial correlation against three types of null models: (i) spatial autocorrelation-preserving spin null models, (ii) degree-preserving rewired null models, and (iii) degree- and edge length-preserving rewired null models (**Fig. 2C**). The observed correlation remained statistically significant across all comparisons, except for the most constrained null model that simultaneously preserved both degree distribution and edge length (*P*_spin_ < 0.005, *P*_rewired1_ < 0.001 and *P*_rewired2_ = 0.058). Moreover, this significant spatial correlation was well replicated in the SUDMEX-CONN cohort (**<u>Fig. S1</u>**). When spatial correlations at the system level were assessed, we found that the propagation of CUD cortical atrophy is mainly constrained by WM connections within the heteromodal association and visual areas (**Fig. 2D**).

**Fig. 2.**
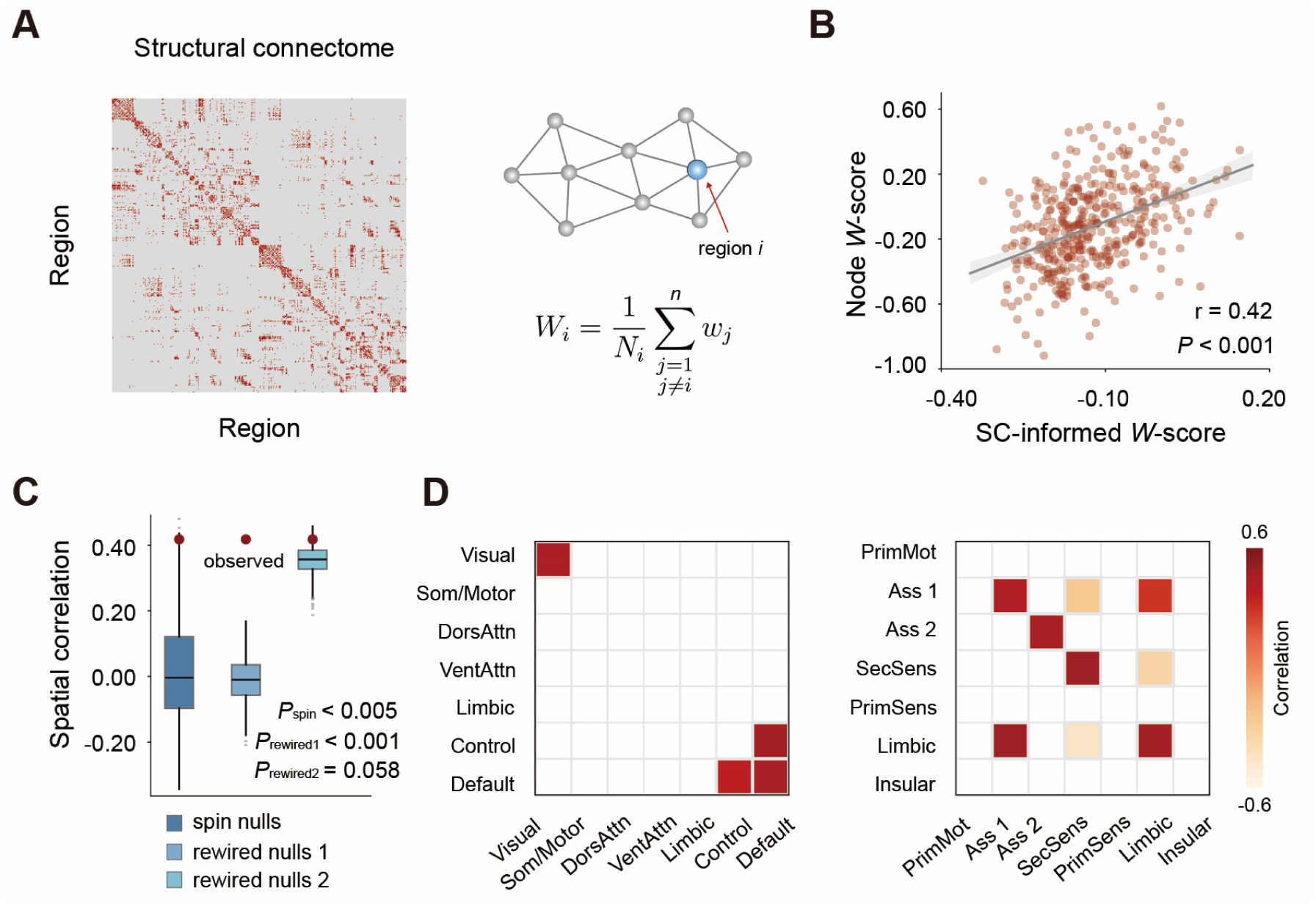
Structural connectome shapes cortical atrophy in CUD. (A) Normative structural connectome backbone constructed using the Schaefer400 atlas^37^. The structural connectome-informed neighbour-level *W*-score for a given region was modelled as the average *W*-score of its structurally connected neighbour regions. (B) Cortical atrophy across the 400 regions was significantly correlated with the mean atrophy of structurally connected neighbours (r = 0.42, *P*_spin/rewired1_ < 0.01 and *P*_rewired2_ = 0.058, one-sided). (C) The observed Pearson’s correlation was compared against three baseline null models. The spin null models were constructed by randomly rotating cortical *W*-score maps on the fs_LR spherical surface, and preserving spatial autocorrelation while randomizing the cortical features (1,000 iterations). The first rewired null model was constructed by randomly rewiring inter-region edges while preserving the degree distribution of the empirical structural connectome (1,000 iterations). The second rewired null model similarly involved random edge rewiring, but preserved the node density, degree distribution and edge length of the empirical structural connectome (1,000 iterations). (D) Region-to-neighbour spatial correlations at the system level. The statistically significant correlations were shown in red (*P*_FDR_ < 0.05, two-sided). SC, structural connectome.

Furthermore, recent findings have shown that disease propagation is related to haemodynamic activity and microscale features, especially the gene expression and neurotransmitter receptor density^24^. Integrating the structural connectome with the biological characteristics of neuronal populations improves the accuracy of propagation pattern prediction models^20^. We thus evaluated whether functional connectivity and molecular feature similarity networks of the brain could account for abnormal cortical thickness patterns in CUD. We masked functional connectivity, transcriptomic similarity and receptor similarity networks using the structural connectome, and used the connection weights derived from these networks to compute the weighted average *W*-score of all structurally connected neighbours for each region (see details in Methods). Greater significant positive correlations were observed between region and neighbour *W*-score values, informed by functional connectivity (**Fig. 3A;** r = 0.53, *P*_spin/rewired2_ < 0.05 and *P*_rewired1_ < 0.001), transcriptomic similarity (**Fig. 3B;** r = 0.57, *P*_spin_ < 0.01 and all *P*_rewired_ < 0.001) and receptor similarity (**Fig. 3C;** r = 0.58, *P*_spin_ < 0.01 and all *P*_rewired_ < 0.001). Additionally, all the results were consistently replicated in the SUDMEX-CONN cohort (**<u>Fig. S1</u>**). This finding extends the brain homophilic mixing theory—that is, connected regions tend to share more similar biological features than unconnected regions^36^—and are also more likely to experience similar atrophy patterns in CUD. Taken together, these results suggest that brain’s structural connectome and local biological properties jointly contribute to the cortical pathology spread in CUD, and highlight the value of annotating WM pathways with local molecular features.

**Fig. 3.**
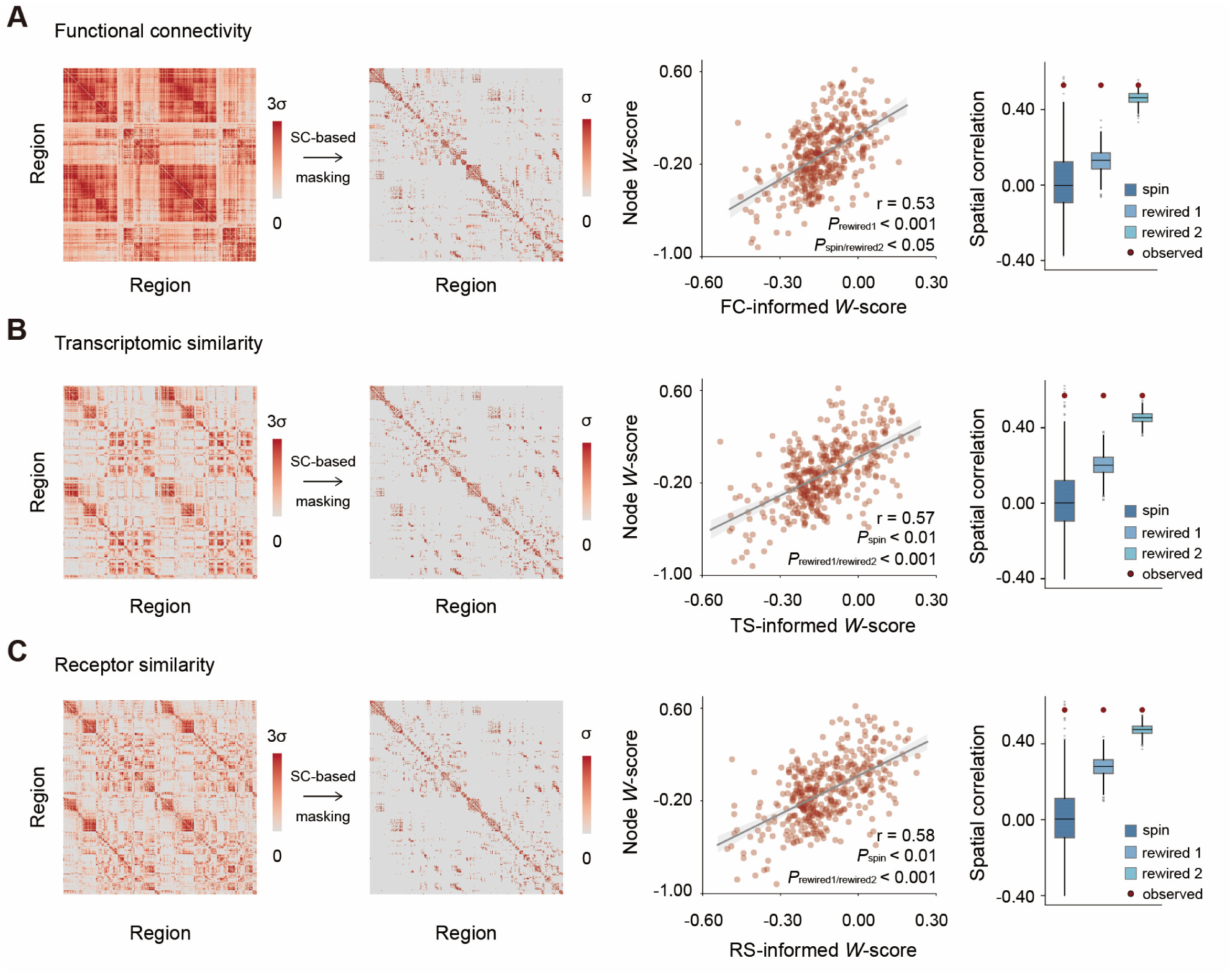
Biological similarity matrices shape cortical atrophy patterns in CUD along the white matter (WM) pathways. (A-C) The inter-regional similarity networks (retaining only positive connections) were first masked by the binary structural connectome and subsequently used as edge weights to calculate the region-to-neighbour spatial correlations, respectively. The atrophy of each region was significantly correlated with the mean atrophy of its structurally connected neighbours, weighted by functional connectivity (r = 0.52, *P*_spin/rewired2_ < 0.05, *P*_rewired1_ < 0.001, one-sided), transcriptomic similarity (r = 0.57, *P*_spin_ < 0.01, all *P*_rewired_ < 0.001, one-sided) and receptor similarity (r = 0.58, *P*_spin_ < 0.01, all *P*_rewired_ < 0.001, one-sided). These inter-regional biological feature similarities consistently amplify the extent of pathology spread compared to structural connectome alone. The significance of correlations was evaluated against the spin and rewired null models (1,000 iterations). FC, functional connectivity; TS, transcriptomic similarity; RS, receptor similarity.

### Multimodal and multiscale connectivity informs CUD epicenters

Given that multimodal and multiscale brain connectivity shapes the spatial distribution of cortical atrophy, we next sought to identify potential epicenters of morphological abnormalities in CUD. Specifically, we define the global disease exposure of region *i* as the average of the products between the edge strength (calculated as *FC_ij_*×*TS_ij_*×*RS_ij_*) and the *W*-score of region *j*, across all regions *j* that are structurally connected to region *i*. Brain regions with both great atrophy and global disease exposure were further identified as epicenters, which not only exhibit disease vulnerability but also facilitate the spread of pathology (**Fig. 4A**)^23,27^. The results revealed that the likeliest disease epicenters were primarily located in the dorsolateral prefrontal cortex (DLPFC), visual cortex, and medial prefrontal cortex (mPFC), and exhibited significant spatial similarity across the two datasets (**Fig. 4B, C <u>and Fig. S2</u>**; r = 0.53, *P*_spin_ < 0.001). Moreover, Stubbs et al.^38^ reported that in the majority of neuroimaging studies on substance use disorders (91% of 45 studies), the reported coordinates of regional brain abnormalities share functional connectivity to a common brain network, referred to as the SUD network. To further validate our results, we assessed the spatial correspondence between the identified CUD epicenters and the SUD network. We retained the top 25% of regions with the highest epicenter likelihood and found that they were part of the SUD network, showing greater overlap than expected under null distributions generated by randomly rotating these “CUD hub” regions (**<u>Fig. S3;</u>** *P*_perm_ = 0.060).

**Fig. 4.**
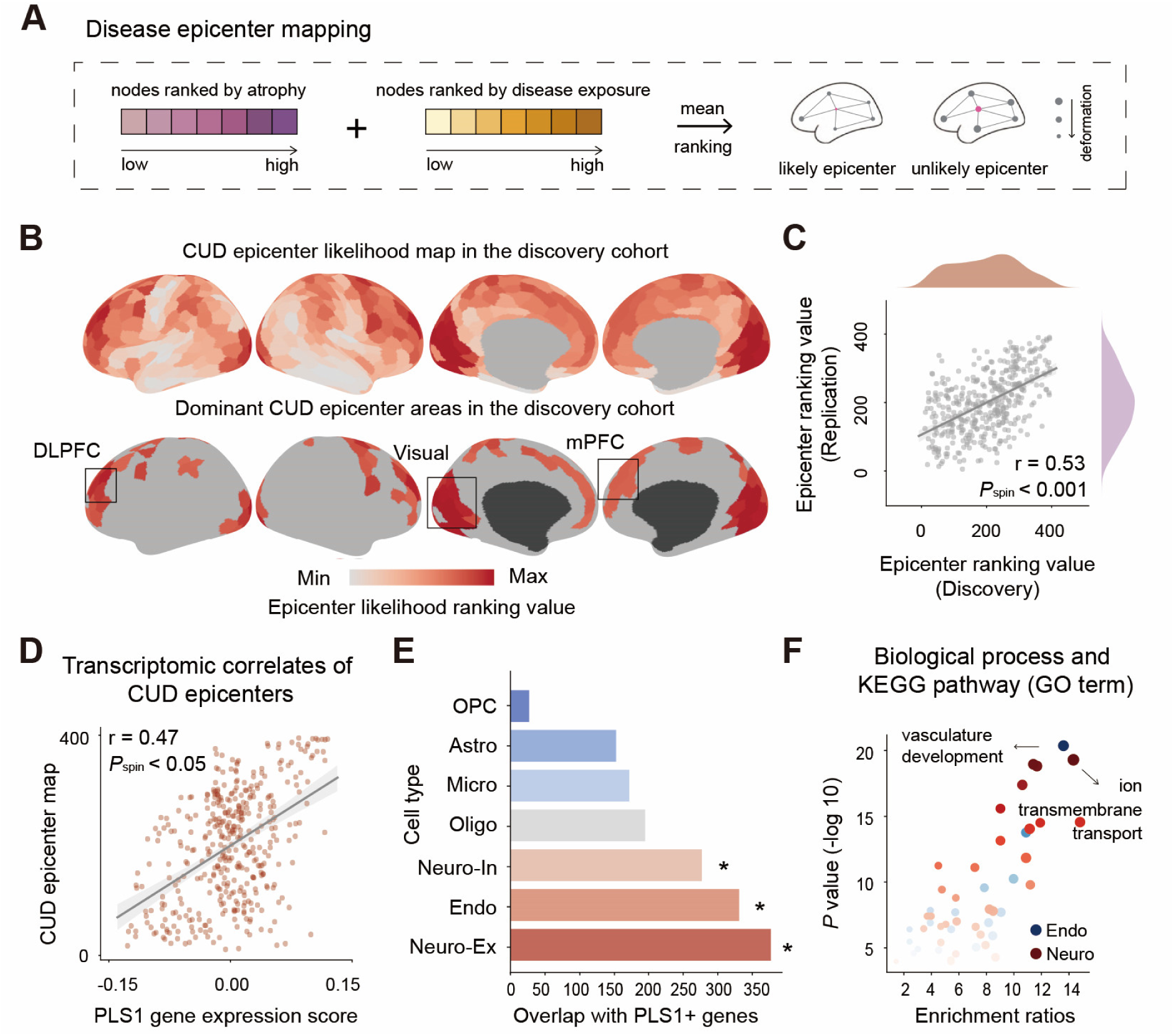
Multimodal and multiscale connectivity-based epicenter identification. (A) The disease exposure of region *i* was defined as the mean of the products between the edge strength (FC*_ij_*×TS*_ij_*×RS*_ij_*) and the *W*-score of each structurally connected region *j*. We hypothesized that if a cortical region exhibits both high atrophy and high disease exposure, it could potentially serve as a pathological epicenter. Regions were ranked based on their atrophy values and disease exposure values, and the average of the two ranks was used to represent the probability of a region being an epicenter. (B) The spatial distribution of CUD epicenter likelihood rankings is shown in the upper panel. The regions with the top 25% of epicenter likelihood rankings were depicted in the bottom panel and were primarily located in the prefrontal and visual cortices. (C) The CUD epicenter likelihood maps were reproducible across the two independent cohorts (r = 0.53, *P*_spin_ < 0.001, one-sided). (D) The epicenter likelihood map was significantly correlated (r = 0.47, *P*_spin_ < 0.05, one-sided) with the primary PLS component (PLS1), which represents a weighted map of 15,632 gene expression scores. (E) The number of overlapping genes between the PLS1+ gene list and cell-type specific genes. All permutated *P* values were FDR-corrected and determined by one-sided tests. Asterisks represent significance at FDR-corrected *P*_perm_ < 0.01. (F) Enriched Gene Ontology (GO) terms of the CUD epicenter-related genes for each cell type. The size of each dot denotes the number of genes within a given term. The transparency of the dot reflects the significance of the corresponding term and the multi-test FDR-corrected *P* values are plotted as log10-transformed values. DLPFC, dorsolateral prefrontal cortex; mPFC, medial prefrontal cortex; OPC, oligodendrocyte precursor; Astro, astrocyte; Micro, microglia; Oligo, oligodendrocyte; Neuro-In, inhibitory neurons; Endo, endothelial; Neuro-Ex, excitatory neurons; KEGG, Kyoto Encyclopedia of Genes and Genomes.

Having established that cortical atrophy in CUD is centered on a set of epicenters, we next asked whether these epicenters are enriched for specific transcriptional signatures. We obtained brain-wide gene expression data from the Allen Human Brain Atlas (AHBA) and parcellated them into region-level resolution (400 regions × 15,632 gene expression data). We used partial least squares (PLS) regression to identify multivariate associations between the regional gene expression metrics and the CUD epicenter likelihood maps. The first PLS component (PLS1) significantly explained 21.84% of the epicenter-transcription covariance (*P*_perm_ < 0.001). A spatial correlation analysis confirmed that the PLS1 weighted gene expression map significantly aligned with the epicenter distribution (**Fig. 4D**; r = 0.47, *P*_spin_ < 0.05), suggesting that the CUD epicenters are embedded within a specific transcriptional landscape. We then ranked the PLS1 genes by their normalized weights and defined the PLS1+ and PLS1-gene sets as those with *Z*-scores > 2 and < –2, yielding 4,802 and 4,204 genes (all *P*_FDR_ < 0.05)^28^. Positively (or negatively) weighted genes were preferentially expressed in regions with greater (or less) epicenter probability, respectively. To further interpret the biological relevance of the epicenter-associated genes, we primarily considered the PLS1+ genes and assigned them to seven neuronal and glial cell types using the approach proposed by Seidlitz et al.^30^. We found that a large part of PLS1+ genes were significantly involved in excitatory neurons (**Fig. 4E**; n = 377, *P*_Perm_ < 0.005), endothelial (n = 331, *P*_Perm_ < 0.005) and inhibitory neurons (n = 277, *P*_Perm_ < 0.005). Enrichment analysis using cell-type-specific genes revealed that neuronal cells associated with CUD epicenters were significantly enriched for Gene Ontology (GO) terms such as “inorganic ion transmembrane transport”, “synaptic signaling”, “modulation of chemical synaptic transmission” and “neuronal projection development” (**Fig. 4F <u>and Supplementary Data 1</u>**). In addition, endothelial cells were significantly enriched for GO terms including “vasculature development” and “circulatory system process”. Collectively, these findings link CUD epicenter-related gene expression to specific cell types and biological processes involved in neuronal communication, synaptic function and neurovascular homeostasis, and highlight the local molecular and cellular features underlying CUD pathology.

### Epicenters associate with clinical responses and psychopathology in CUD

Next, we explored whether the multimodal and multiscale connectivity-informed epicenters could provide insight into intervention strategies for CUD. We used left DLPFC repeated transcranial magnetic stimulation (rTMS) coordinates **<u>(Fig. S4)</u>**, functional connectivity data, and symptom-specific behavioural improvements for each patient (N = 15, age range: 25-47 years, 2 females) to compute distinct symptom-response cortical maps^39^. These response maps quantify the extent to which each voxel’s functional connectivity to the stimulation site predicts treatment efficacy for a given CUD symptom. Within the maps, positive values indicate that the region was more positively correlated with effective rTMS sites, whereas negative values indicate stronger negative correlations with those sites associated with higher clinical efficacy. We found that the identified epicenter likelihood map was negatively correlated with the Visual Analog Scale (VAS)-response map (**Fig. 5A**; *t*_396_ = −4.50, *P*_FDR_ < 0.001) and Cocaine Craving Questionnaire NOW (CCQN)-response map (*t*_396_ = −3.91, *P*_FDR_ < 0.001). These findings suggest that regions more likely to be CUD epicenters are more strongly negatively correlated with the effective rTMS site for reducing cocaine craving, and highlight the relationships between CUD epicenters and neuromodulation mechanisms.

**Fig. 5.**
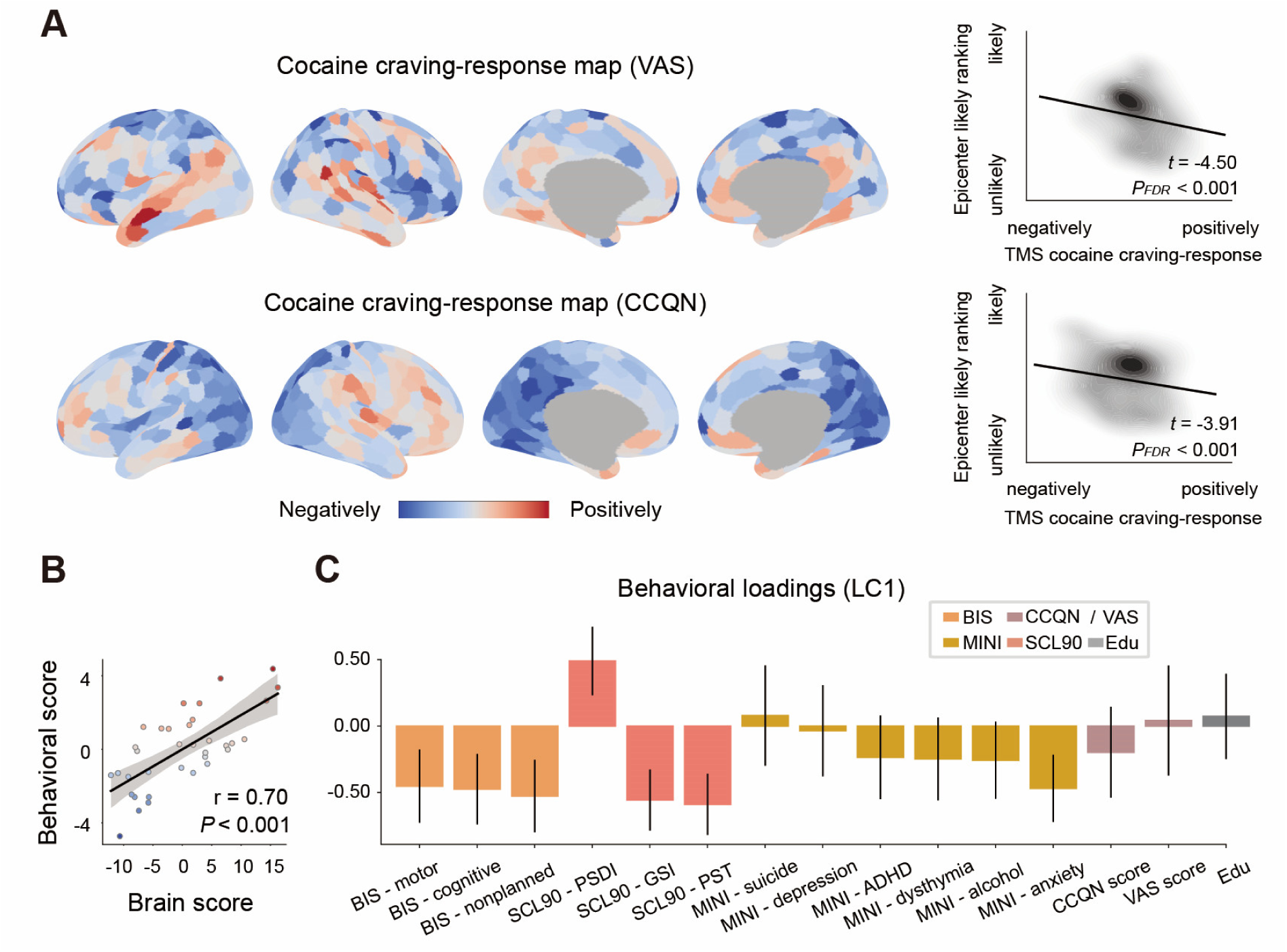
CUD epicenters correlate with clinical responses to neuromodulation and psychopathology severity. (A) Repeated transcranial magnetic stimulation (rTMS) response map for the Cocaine Craving Visual Analog Scale (VAS) and the Cocaine Craving Questionnaire Now (CCQN), two clinical metrics used to assess cocaine craving status (left panel). The epicenter likelihood map was significantly correlated with the VAS response map (*t*_396_ = −4.50, *P*_FDR_ < 0.01, two-sided) and the CCQN response map (*t*_396_ = −3.91, *P*_FDR_ < 0.01, two-sided) estimated from the SUDMEX-TMS cohort. These *t*-values were obtained from a multivariate linear regression model using ordinary least squares, in which epicenter likelihood values were predicted by VAS and CCQN response maps. (B) A significant correlation between individuals’ brain scores and behavioural scores on the first latent component (LC1) derived from a partial least squares (PLS) analysis (r = 0.70, *P* < 0.001). Each scatter point represents an individual. (C) Behavioural loadings of the LC1. Greater loading on LC1 was associated with milder psychopathological manifestations across multiple domains, including fewer psychiatric symptoms and lower levels of cocaine craving. Error bars indicate the standard deviation of bootstrap resampling (1,000 iterations). BIS, Barratt Impulsiveness Scale; SCL90, Symptom Checklist-90-revised; PSDI, Positive Symptom Distress Index; GSI, Global Severity Index; PST, Positive Symptoms Total; MINI, Mini International Neuropsychiatric Interview; ADHD, attention-deficit hyperactivity disorder; Edu, education.

Finally, to determine whether CUD cortical epicenters could reflect individual variability in clinical behaviour assessments, we examined the covariance between individual epicenter likelihood maps and individual clinical profiles using PLS analysis. Because age and sex were controlled in the atrophy modelling process, they were also regressed from the behavioural data. The first latent component (LC1) was statistically significant (*P*_Perm_ < 0.001), accounting for 56.07% of the total covariance, and could be cross-validated (**<u>Fig. S5;</u>** 2-fold, *P* < 0.05). The LC1 captured a spatial pattern of epicenter variation, with the highest loadings in the lateral temporal cortex, insular and frontal opercular cortex **<u>(Fig. S6)</u>**, and exhibited a significant association between the brain and composite behavioural scores (**Fig. 5B;** r = 0.70, *P* < 0.001). Patients with higher expression of this epicenter pattern tend to exhibit more over-controlled personality traits (BIS scores), lower psychological distress (SCL90 scores), fewer psychiatric symptoms (MINI scores), and reduced cocaine craving (CCQN score; **Fig. 5C**). Together, these results suggest that LC1 may reflect the general psychopathology of CUD, and that individual differences in the epicenter pattern are closely associated with the symptom severity and behavioural characteristics of CUD patients.

### Results were robust across a range of sensitivity analyses

The study comprised a series of sensitivity analyses to ensure the robustness of the major findings. First, we demonstrated that the associations between regional *W*-scores and the *W*-scores of structurally-connected neighbours were also significant in an independent replication dataset. These correlations were further strengthened when accounting for inter-regional similarity in hemodynamic activity and molecular features (**<u>Fig. S1</u>**). Second, we adopted a Susceptible-Infected-Removed (SIR) agent-based model to identify the CUD epicenters. This model simulates local vulnerability modulated by protein misfolding in the cortex and the trans-neuronal spreading process of pathological proteins observed in diseases. We found that the epicenter likelihood maps derived from the data-driven ranking-based method and the SIR method exhibited significantly correlated spatial distribution patterns (**<u>Fig. S7</u>**; r = 0.66, *P*_spin_ < 0.001). Third, the principal component of the behavioural measures derived from principal component analysis (PCA) showed a highly positive correlation with the behavioural scores of LC1 (r = 0.53, *P* < 0.001). Moreover, instead of computing covariance to represent the importance of each behavioural measure for LC1, we employed Pearson’s correlation to assess this importance. The results showed that the behavioural loadings on LC1 obtained from the two analytical approaches were highly consistent (**<u>Fig. S8;</u>** r = 0.95, *P* < 0.001).

## Discussion

This work presents a structural connectome- and regional biological attribute-based pathology spreading framework for CUD, and identifies neuroimaging biomarkers to bridge neural and clinical symptom profiles. Specifically, we found that the atrophy patterns of cortical regions are significantly correlated with the mean atrophy of structurally-connected neighbouring regions. Moreover, integrating inter-regional similarity in biological features can improve the inference of the spatial patterning of atrophy in CUD, providing insight into the molecular mechanisms underlying neuroimaging phenotypes. By mapping multimodal and multiscale connectivity onto the cortical atrophy patterns in CUD, we identified CUD-related epicenters. Cortical areas with high epicenter likelihood rankings are primarily located in the prefrontal and visual cortices, and are associated with genes enriched for biological pathways involved in neuronal communication, synaptic function and neural homeostasis. Finally, the spatial distribution of epicenters shows potential clinical relevance and corresponds to individual behavioural symptoms. Our findings are replicated across two independent datasets and a series of sensitivity analyses. Taken together, these findings provide insights into the network-level mechanisms underlying cortical atrophy in CUD and establish potential neural targets for clinical intervention.

### Normative modelling-derived cortical atrophy in CUD

CUD has been characterized by widespread structural brain abnormalities in numerous neuroimaging studies. Although cortical thinning and gray matter reduction have been consistently observed, the findings across these studies are heterogeneous and sometimes contradictory. For example, Bittencourt et al. and Makris et al. consistently reported significant cortical thinning in the left orbitofrontal cortex^12,40^, whereas Mackey et al. found no evidence of structural alterations in the prefrontal cortex and instead reported a significantly thinner supramarginal cortex^13^. Indeed, such population-level differences obscure the heterogeneity across individuals and may not be representative of individual cases. Using normative modelling, we estimated the normative range of variation for cortical thickness and quantified person-specific deviations from the healthy population expectations for each patient with CUD. Compared with previous studies, our results consistently characterized cortical thinning in the frontal cortices, which has been associated with cognition changes^41^ and years of cocaine dependence^40^. Additional complementary evidence also supports the notion that brain structure abnormalities in the frontal cortex are a consistent feature of CUD, including reduced gray matter volume^10^ and lower gray matter tissue density^42^. CUD also shares similar cortical atrophy patterns in the frontal cortex with other psychiatric disorders^32^, indicating the overlap of neural substrates. Moreover, our results further revealed significant cortical atrophy in the visual cortex, whose dysfunction has recently been implicated in the functional circuit disruption of CUD^6,43^. At the micro level, these multifaced gray matter changes are underpinned by multiple biological processes, such as synaptic pruning^44^ and neurofibrillary tangles^45^. Building a unified etiologic model to account for the brain-wide morphological changes in CUD is challenging, as atrophy in different regions may have distinct biological origins. We thus attempt to address this issue through network-based modelling approaches.

### Pathology spread hypotheses in CUD

The principal finding of the current study is that regional atrophy patterns can be well represented by those of structurally connected neighbours in CUD. Moreover, atrophy patterns are conditioned not only by the WM structural network but also by multiple biological similarity networks. These findings were highly reproducible across two independent datasets. The observed network-based atrophy could be explained by several underlying mechanisms. **<u>First</u>**, cortical regions that are connected by WM pathways are likely to share similar cytoarchitectural and morphological properties^46,47^, including cellular size, neuronal density, and myelination^48,49^, all of which are closely associated with gray matter alterations. **<u>Second</u>**, direct WM connections facilitate interregional communication; therefore, these regions tend to exhibit synchronous spontaneous neural activity^50,51^ and coordinated changes in brain morphometric features^52,53^. **<u>Third</u>**, both cocaine-induced structure changes and architecture of WM connectome are highly relevant with genetic factors and gene expression alterations^54–56^. This molecular-level mechanism may involve a synergistic regulatory process, which manifests at the macroscopic level as propagating patterns of morphological atrophy along the structural connectome. **<u>Fourth</u>**, cortical regions that are structurally connected are also likely to share similar neurotransmitter receptor profiles^57^, and neurotransmitter systems are closely related to gray matter changes in CUD. For example, cocaine acutely targets the human dopamine system, and dopamine accumulation increases metabolic activity, subsequently leading to axon terminal degeneration, neuronal death, and receptor downregulation, which may trigger a pattern of atrophy that spreads along the structural connectome and is shaped by underlying transcriptomic and receptor similarities^41^. Together, the present findings suggest a pathology-spreading framework in which the course and expression of CUD are mediated by both the structural connectome and local biological vulnerability. Specifically, local features, including intrinsic neural activity, gene expression and receptor distribution, contribute to regional selective vulnerability, whereas pathological processes spread through WM pathways, guided and amplified by inter-regional similarity in biological attributes.

### Disease epicenters in CUD

Given that atrophy and brain connectivity are related in CUD, we next developed a multimodal, multiscale connectivity-based node ranking approach to examine the regional vulnerability. By mapping multimodal and multiscale connectivity profiles onto cortical atrophy patterns in CUD, we identified regions in the DLPFC, visual area, and mPFC emerged as the most likely disease epicenters, whose connectivity profiles spatially resembled the observed CUD-related atrophy patterns. **<u>These</u> <u>prefrontal cortical regions</u>**, primarily situated within the frontoparietal control and default mode networks, have been implicated in atypical changes associated with substance addiction, including patterns of resting-state functional connectivity^6,43^, functional dynamics^58^, structural covariance features^26^, transcriptional alterations^59^, and structural connectivity^60^. Dysfunction of the prefrontal cortex and its associated circuity has been strongly linked to the impaired inhibitory control in CUD^61–64^. Within the prefrontal cortex, DLPFC is known to mediate top-down regulation of negative affect, while mPFC is involved in the monitoring and modulation of affective states^65,66^. Therefore, these regions are also frequently implicated in psychiatric symptoms across a wide range of disorders, including depression^67,68^, schizophrenia^69,70^ and risky decision-making^71–73^. **<u>As for the visual regions</u>**, recent data-driven studies have increasingly expanded the focus of psychiatric illnesses beyond higher-order association cortex and highlighted the role of occipital regions as well as sensorimotor cortices^74–77^. For example, Zhao et al. reported increased resting-state functional connectivity between the visual network and dorsal attention network in CUD, along with significant associations between visual-dorsal attention connections and impulsivity traits^6^. Higher transdiagnostic general scores (that is, the *p* factor) are associated with hyperconnectivity between the visual association cortex and both frontoparietal control and default mode networks^76^. Taylor et al. identified a transdiagnostic brain network for psychiatric illness defined by positive functional connectivity to the anterior cingulate and insula, and negative connectivity to the posterior parietal and lateral occipital cortex^78^. Consistent with these findings, Stubbs et al. further reported a common brain network for substance use disorders characterized by negative connectivity to the medial prefrontal and occipital cortices^38^. Together, our results suggest that abnormalities in the DLPFC, visual cortex, and mPFC are likely causes or consequences of CUD.

What microscale features predispose the prefrontal and visual regions to act as macroscale hubs in CUD? We identified a positive correlation between the epicenter likelihood map and the weighted cortical expression of genes, which primarily enriched for neuronal communication, synaptic function and neurovascular homeostasis. These results are in line with the theory that cocaine inhibits the reuptake of multiple neurotransmitters (in particular dopamine and serotonin) into presynaptic neurons by blocking their respective transporters^56^, and that chronic cocaine use induces synaptic plasticity, such as enhanced synaptic transmission between dopamine neurons^79^. In addition, the distribution of CUD epicenters significantly overlaps with the transcriptomic signatures of endothelial cells, which form the blood‒brain barrier (BBB) and contribute to maintain the homeostasis of the brain microenvironment^80–82^. Indeed, multiple rodent studies have provided evidence for the cocaine-induced breach in BBB^83,84^. Together, these findings reveal the transcriptomic and cellular underpinnings of regional brain vulnerability in CUD.

Furthermore, using an approach developed by Siddiqi et al.^39^, we assessed the group-level therapeutic response of CUD patients to left DLPFC rTMS across the cortex. We found that the multimodal and multiscale connectivity-informed epicenters were spatially correlated with the cocaine craving-response maps. This finding suggests that symptom improvements in cocaine addiction among CUD patients can be effectively elicited by stimulation of the major epicenter, that is, the DLPFC. This result is consistent with prior studies showing that lesions causing remission from psychiatric disorders intersect with brain networks derived from atrophy maps in patients^38,78^. We also demonstrated that individual-specific epicenter distributions in CUD patients account for individual variations in multivariate clinical manifestations, including cocaine craving and the co-occurring symptoms such as impulsivity, attention deficit and anxiety. Together, given that the therapeutic outcomes of rTMS are closely related to the connectivity profiles of stimulation targets^85–88^, and that the underlying connectivity of epicenters constrains CUD-related cortical abnormalities, our findings provide the rationale for the therapeutic effects of brain stimulation and inform the potential rTMS targets in CUD treatment.

### Limitations

Several methodological issues merit further consideration. First, the cross-sectional nature of this study limits the ability to investigate how CUD pathology evolves over time. Future studies incorporating longitudinal data could help elucidate the effects of disease progression on cortical atrophy patterns and putative epicenters. Second, we used a structural connectome derived from the healthy population as a normative representation of the brain’s wiring diagram and demonstrated that the normative connectome and cortical atrophy are related in CUD, supporting a network-based pathology spreading framework. To characterize the connectome-based architectural foundation of brain morphology accurately, our approach did not account for CUD-related abnormalities in white matter tracts, functional connectivity, transcriptomic similarity, or receptor similarity across regions. As a result, our analyses are correlational, which limits our ability to draw causal inferences. For instance, alternative possibilities warranting further investigation include: (1) white matter tract abnormalities may emerge first, disrupting inter-neuronal communication and the transport of trophic factors, thereby contributing to widespread cortical atrophy; or (2) localized morphological changes may originate in vulnerable regions and subsequently propagate, ultimately altering inter-regional structural and functional connectivity. Modeling the causal directionality of pathological processes in CUD represents a promising and important avenue for future research. Finally, although we demonstrated the clinical relevance of CUD epicenters through correlations with pre-existing rTMS data, future studies are needed to explore the feasibility of directly applying individual-specific epicenter likelihood maps to guide targeted neuromodulation.

### Conclusions

In conclusion, we provide evidence that CUD-related cortical atrophy reflects the brain’s structural connectome, functional connectivity and molecular architecture. Aligning CUD-related cortical changes to a reference frame of multimodal connectome and multiscale biological attributes allows us to systematically conceptualize the pathological processes underlying CUD, thereby helping to identify effective therapeutic targets.

## Methods

### Discovery cohort

#### Participants and MRI acquisition

The SUDMEX-TMS dataset was used as the discovery cohort in this study^89^. This dataset includes 54 participants, who were recruited in Mexico City, Mexico, and met the criteria of having cocaine dependence for at least 12 months, with a frequency of at least 3 days per week and no more than 60 continued days of abstinence in the past year. All participants provided written informed consent, and all procedures were approved by the Institutional Ethics Research Committee (CEI/C/070/2016) and registered on ClinicalTrials.gov (NCT02986438). The neuroimaging data of one patient was discarded by the data acquisition team, resulting in a final sample of 53 individuals with CUD (age range:18-48 years, 8 females) included in the subsequent analyses.

Anatomical T1-weighted (T1w) images were acquired using a 3D FFE SENSE sequence on a Philips Ingenia 3T MR system with 32-channel heal coil. The acquisition parameters for the structural images were as follows: repetition time / echo time = 7 / 3.5 ms, field of view = 240 mm^2^, matrix = 240 × 240 mm, slices = 180, gap = 0, plane = sagittal, voxel = 1 × 1 × 1 mm. Eyes-open resting state functional MRI (fMRI) data were acquired using a gradient recalled echo planar imaging sequence with the parameters as follows: repetition time / echo time = 2000 / 30 ms, flip angle = 75°, matrix = 80 × 80, field of view = 240 mm^2^, voxel size = 3 × 3 × 3.33 mm, number of slices = 36, slice acquisition order = interleaved, and phase encoding direction = anterior-to-posterior.

#### Clinical assessments

Among the CUD patients, cocaine dependence was diagnosed using the Mini International Neuropsychiatric Interview-Plus (MINI) Spanish version 5.0.0. Diagnoses for anxiety, alcohol use, attention-deficit, depression, dysthymia and suicide risk of patients were also assessed via the MINI. Impulsive behaviours were evaluated using the Barratt Impulsiveness Scale version 11 (BIS-11) including three domains: cognitive impulsiveness, motor impulsiveness and non-planning impulsiveness. The Symptom Checklist-90-revised (SCL-90) was used to assess the psychological symptoms and distress in 3global measures: Global Severity Index (GSI), Positive Symptom Distress Index (PSDI) and Positive Symptoms Total (PST). Cocaine craving was assessed by both the CCQN and the VAS scale.

#### rTMS treatment

Across the 53 included CUD patients, 8 discontinued participation in the rTMS intervention, resulting in 25 patients completing the 2-week acute rTMS treatment in the active group and 20 in the sham group, following a double-blind, randomized controlled trial^89^. A MagPro R301 Option magnetic stimulator and a figure-of-eight B65-A/P coil were used for stimulation. The acute phase consisted of 10 weekdays of treatment, with 5,000 pulses delivered per day. Specifically, each day included two sessions, each comprising 50 trains at 5 Hz (50 pulses per train), with a 10-second inter-train interval and a 10-minute inter-session interval. Stimulation was delivered to the left DLPFC at 100% of the motor threshold. MRI scans and clinical assessments were conducted both before and after the acute phase; however, post-treatment neuroimaging data were not analyzed in the present study. Although the SUDMEX-TMS cohort also included a maintenance phase, it was not considered in this study because of the limited sample size.

### Replication cohort

#### Participants and MRI acquisition

We used the SUDMEX-CONN dataset as a replication dataset in the present study, which originally recruited 74 patients with CUD and 64 healthy controls^90^. Two healthy controls were excluded due to missing demographic information, resulting in a final sample of 74 CUD patients (age range:18-50 years, 9 females) and 62 healthy controls (age range:18-48 years, 11 females) for replication analyses. The diagnostic criteria for identifying CUD patients were consistent with those used in the discovery cohort, and all participants provided written informed consent. T1w data were scanned using a three-dimensional FFE SENSE sequence on a Philips Ingenia 3T MR scanner with a 32-channel ds Head coil, using the same acquisition parameters as those applied in the SUDMEX-TMS cohort. The SUDMEX-CONN study was approved by the ethics committee of the Instituto Nacional de Psiquiatría “Ramón de la Fuente Muñiz”.

### HCP cohort

#### Participants and MRI acquisition

Our study also included 302 non-substance use individuals (age range: 22-36 years, 219 females) for normative modelling, and 326 unrelated healthy individuals (age range:22-35 years, 145 females) for normative brain connectome construction, all obtained from the Human Connectome Project (HCP)-Young Adult (HCP-YA) dataset^91^. Non-substance use individuals were defined as those who did not meet the diagnostic criteria for any substance use disorder, were not binge drinkers, and reported consuming fewer than 2 alcoholic drinks per day on average over the last 12 months^58^. Unrelated participants were selected to eliminate the influence of familial relationships^24,92^. All data were acquired using a customized Siemens 3T Skyra scanner with a 32-channel head coil. For each participant, two separate T1w images were acquired using the 3D MPRAGE sequence and subsequently averaged. The acquisition parameters were as follows: repetition time / echo time = 2400 / 2.14 ms, field of view = 224 mm^2^, matrix = 320 × 320 mm, slices = 256, plane = sagittal, voxel = 0.7 × 0.7 × 0.7 mm. All participants provided informed consent, and the HCP-YA study was approved by the Institutional Review Board at Washington University in St. Louis. Additional details regarding the data collection procedures are available elsewhere^93^.

### MRI data preprocessing

All T1w images from the SUDMEX-TMS and SUDMEX-CONN datasets were preprocessed using FreeSurfer (version 7.2.0), implemented within the fMRIPrep docker container (version 22.1.1)^94^. Specifically, the T1w images were first corrected for intensity non-uniformity and skull-stripped. Brain tissue segmentation was performed using FSL’s FAST, and brain surfaces were reconstructed from the T1w images with FreeSurfer’s recon-all. Finally, space normalization to standard space (MNI152NLin6Asym) was performed via nonlinear registration with antsRegistration, using brain-extracted versions of both the T1w and the standard template. The Scharfer400 parcellation was transformed into the T1w surface space of each participant, and cortical thickness was measured across 400 gray matter cortical regions. Additionally, preprocessed T1w images from the HCP-YA cohort were publicly available and provided by the HCP consortium. The preprocessed steps include bias field correction, space normalization, segmentation and surface reconstruction. The details of the HCP minimal preprocessing pipeline can be found elsewhere^95^.

Resting-state fMRI data from the SUDMEX-TMS cohort were preprocessed using the fMRIPrep pipeline (version 22.1.1)^94^. Specifically, the preprocessing steps include slice timing correction, susceptibility distortion correction, co-registration to T1-weighted anatomical images using boundary-based registration, and spatial normalization to standard MNI space. Confound regressors—including motion parameters, global signals, aCompCor/tCompCor components, and motion outliers (framewise displacement > 0.5 mm or DVARS > 1.5)—were computed. Additional denoising was then performed using ICA-AROMA following spatial smoothing (6 mm FWHM).

### Normative modelling for cortical thickness and CUD deviations

A general linear model was trained with data (N = 364) from the HCP-YA cohort and healthy controls from the replication cohort to estimate normative expectations for regional cortical thickness variations, with age and sex effects statistically controlled^96^. The beta terms of all covariates, along with the residual terms, were then used to calculate the patient-specific cortical thickness deviations in relation to the trained normative model. A *W*-score was computed for each region of each patient using the following formula:

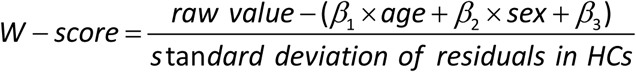

where all beta values were derived from the normative model trained on the healthy controls’ data.

### Normative brain connectome construction

Multimodal imaging data (N = 326), acquired as part of the HCP-YA cohort and described in detail in Hansen et al.^24^, were used to construct the high-quality brain connectome in healthy population. Briefly, diffusion MRI data were processed using MRtrix3^97^, including constrained spherical deconvolution, probabilistic tractography, and SIFT2-based filtering, to reconstruct and refine WM streamlines. The estimated streamlines were then projected to the Schaefer400 parcellation to generate the individual structural connectivity matrix. A group-consensus binary structural connectome was constructed using a density- and edge length-preserving approach that retains edges based on their frequency across subjects within distance-based bins. Preprocessed eyes-open regional blood oxygen level dependent (BOLD) signals were extracted based on the Schaefer400 parcellation, and functional connectivity across regions was calculated as the Pearson correlation coefficient between the regional BOLD time series. A group-average functional connectome was estimated as the mean of pairwise connections across four runs and individuals, and the resulting correlation coefficients were Fisher’s z-transformed. Finally, negative connections were thresholded (i.e., set to zero) to rule out potential confounding effects^23^.

### Normative molecular similarity network construction

The transcription data used to construct the gene expression similarity network are provided by AHBA (https://human.brain-map.org). The microarray expression data were collected from 6 postmortem brains (age range: 24-57 years, 1 female), processed with the abagen toolbox and mapped on the Schaefer400 parcellation, resulting in a 400×8,687 regional gene expression matrix for each donor. The processed steps within the abagen included probe reannotation and filtering, selection of stable probes across donors, spatial mapping of tissue samples to brain regions, normalization of gene expression values, region-wise sample averaging per donor, and filtering genes by stability^24,98^. Furthermore, the regional gene expression matrices were averaged across donors and the normalized gene expression profiles of each region were correlated (Pearson’s r) with each other to construct the transcriptomic similarity network. Finally, negative connections were thresholded (i.e., set to zero) to eliminate potential confounding effects.

To construct the receptor similarity network, positron emission tomography (PET) tracer images were used to measure the receptor density profiles of 18 neurotransmitter receptors/transporters at cortical regions^24,57^. These receptors and transporters span 9 different neurotransmitter systems, including dopamine, norepinephrine, serotonin, acetylcholine, glutamate, gamma-aminobutyric acid (GABA), histamine, cannabinoid, and opioid. PET tracer data were mapped to the Schafer400 parcellation and *Z*-score normalized. The receptor profiles of each region were correlated (Pearson’s r) with each other and the resulting correlation coefficients were Fisher’s z-transformed. Finally, negative connections were thresholded (i.e., set to zero) to eliminate potential confounding effects.

### Neighbourhood atrophy estimation

We used the normative structural connectome to define the neighbours of each cortical region. The average atrophy of structural neighbours for the *i*-th cortical region was calculated as follows:

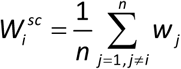

where *n* is the total number of regions that are structurally connected to region *i*, and *w_j_* is the *W*-score of neighbour region *j*. The weighted average atrophy of functional neighbours for the *i*-th cortical region was calculated as follows:

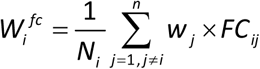

where *N_i_* is the total functional connection strength of region *i*, *n* is the total number of regions that are structurally connected to region *i*, *w_j_* is the *W*-score of neighbour region *j*, and *FC_ij_* is the functional connectivity between region *i and* region *j*. The weighted average atrophy of transcriptomic neighbours and receptomic neighbours for the *i*-th cortical region was calculated in the same manner as above. Finally, Pearson correlation analysis was performed to assess the relationship between regional atrophy and the collective atrophy of their neighbours, a procedure that was repeated four times.

### Epicenter identification

We developed a multimodal and multiscale connectivity-based node ranking approach to identify disease epicenters in individual patients, building on the phenomenon that WM projections provide routes for the spread of CUD pathology and such spreading is more likely to occur between regions with similar biological properties. Specifically, regional atrophy and regional disease exposure were ranked separately^23^. The disease exposure of cortical regions was defined using the brain connectome and molecular architecture as follows:

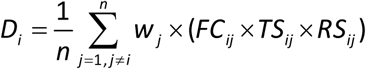

where *n* is the total number of regions that are structurally connected to region *i*, *w_j_* is *W*-score of neighbour region *j*, *FC_ij_* is the functional connectivity between region *i and* region *j*, *TS_ij_* is the correlated transcriptomic similarity between region *i and* region *j* and *RS_ij_* is the correlated receptor similarity between region *i and* region *j*. The mean ranking value was used to represent the probability of a region being an epicenter. A higher value indicates that the region exhibits more severe cortical thinning and is also connected to regions with greater atrophy. Finally, the individual epicenter likelihood maps were averaged across patients to generate the group-level epicenter likelihood map.

### Gene enrichment analysis

We utilized gene expression data from the AHBA dataset (http://human.brain-map.org), which were derived from six postmortem brains with 3702 spatially distinct samples. The detailed preprocessing procedures can be found in the method section ‘Normative molecular similarity network construction’. The only modification was that, to retain more genes for enrichment analysis, we did not apply stability-based gene filtering as described previously. As a result, 15,632 genes remained, and a gene expression matrix (400 × 15,632) was used for subsequent analyses.

PLS regression was employed to identify multivariate relationships between the epicenter likelihood map and the transcriptomic profiles of all 15,632 genes^98^. In the PLS model, Z-normalized gene expression data served as the predictor variable, and Z-normalized epicenter likelihood values were employed as the response variables. PLS1 captures the optimal low-dimensional representation of gene expression profiles that are mostly correlated with the epicenter likelihood map. The statistical significance of the variance explained by PLS1 was assessed by spatially permuting the response variables 1,000 times. Bootstrapping was then applied to quantify the estimation errors in the PLS1 weight of each gene, and *Z* scores were calculated by dividing the weight of each gene by its bootstrap standard error, enabling genes to be ranked based on their contribution to PLS1.

Using published cell-type-specific gene lists from Seidlitz et al.^30^, we categorized the PLS1+ genes (*Z* > 2) into seven canonical classes: excitatory and inhibitory neurons, astrocytes, endothelial cells, microglia, oligodendrocyte precursors and oligodendrocytes. The significance of the overlap between each cell type and the PLS1+ genes was assessed by a permutation test, in which random gene sets of equal size were sampled from the gene pool and their overlap with the PLS1+ genes were calculated across 1,000 iterations, followed by false discovery rate (FDR) correction. Finally, PLS1+ genes involved in each cell type were submitted separately to the Metascape website, an automated functional pathway enrichment tool, to perform enrichment analysis and provide biological insights into the given genes^99^. All identified biological pathways were considered significant at FDR-corrected *P* values less than 0.05.

### rTMS-induced symptom-response mapping

Following a previous study^39^, we performed seed-based connectivity analysis to generate the symptom-response maps of targeted intervention, showing connections with the effective stimulation site that most correlated with the CUD-related symptom improvements. All patients (N=15, age range: 25-47 years, 2 females) from the active group of the SUDMEX-TMS cohort with both stimulation target localization information and functional MRI data were included in this analysis. Specifically, we first computed vertex-wise functional connectivity matrices for each individual using Pearson correlation, and extracted the connectivity of each stimulation site within a 4 mm radius. We then computed the correlation between these stimulation site connectivity maps and the absolute change of each symptom across subjects. This procedure yielded a value at each vertex, and we refer to the resulting cortical maps as symptom-response maps. Finally, we assessed the relationships between the epicenter likelihood map and the cocaine craving response maps using a multiple regression model as follows:

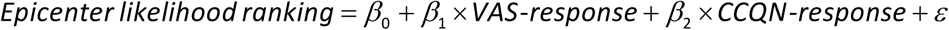

FDR-corrected *P* values less than 0.05 were considered statistically significant.

### Multivariate associations between individual epicenters and behaviors

PLS analysis was performed to examine multivariate associations between individual epicenter likelihood maps and clinical measures. PLS decomposes the cross-covariance matrix between two components into latent components (LCs) that capture the maximal shared variance^100^. Participant-specific brain and behavioural scores for each LC were obtained by projecting the original data onto the corresponding brain and behavioural saliences. The statistical significance of each LC was assessed via 1,000 permutations, and the resulting *P* values were corrected for multiple comparisons via FDR correction.

To interpret the LCs and assess the contributions of individual features, we performed bootstrap resampling (1,000 iterations) by randomly sampling participants with replacement. For each bootstrap sample, the PLS analysis was repeated, yielding a distribution of feature weights across iterations. A bootstrap ratio was calculated for each behavioural feature as the ratio between its original weight to the standard error estimated from the bootstrap distribution. Features with higher bootstrap ratios were considered more strongly associated with the latent variable. Finally, the multivariate association patterns were assessed via repeated 2-fold cross-validation (150 iterations), computing out-of-sample correlations between the brain and behavioural scores. Statistical significance was evaluated via a non-parametric permutation test (150 repetitions) to generate a null distribution of out-of-sample correlations.

### Null models

The observed spatial correlations were compared against three null models. The spin null model preserved spatial autocorrelation while randomizing the cortical features by rotating the feature maps randomly (1,000 iterations). Permutation *P* values were calculated by counting the number of times the null correlation exceeded (for positive correlations) or fell below (for negative correlations) the observed correlation, divided by the total number of spatial rotations. This spin null model was implemented using the Neuromaps toolbox (https://github.com/netneurolab/neuromaps).

The two rewired null models were constructed to evaluate whether the observed correlation was determined by the topology of the connectome itself rather than by basic network properties. In the first rewired model, we used the Maslov-Sneppen algorithm to rewire the structural connectome while preserving the degree distribution of each node (1,000 iterations), implemented via the Brain Connectivity Toolbox (randmio_und_connected.m; https://brainconn.readthedocs.io). In the second rewired model, edges were randomly swapped to rewire the network while preserving the node density, degree distribution and edge length (based on the Euclidean distance), consistent with the empirical structural connectome (1,000 iterations; https://www.brainnetworkslab.com/coderesources). The *P* value was calculated as the proportion of correlations in the null models that exceeded the observed correlation (for positive correlations) or fell below the observed correlation (for negative correlations).

## Author contributions

Z.H., T.L., G.Y. and T.Y. conceptualized and designed the study; Z.H. analyzed the data and wrote the first draft of the manuscript; K.W. contributed to data processing and figure generation; G.Y., T.L. and T.Y. jointly supervised the project and edited the manuscript.

## Data availability

The raw neuroimaging and behavioural data from the SUDMEX-TMS and SUDMEX-CONN datasets are publicly available on OpenNeuro (https://openneuro.org/datasets/ds003037/versions/2.1.0; https://openneuro.org/datasets/ds003346/versions/1.1.2). Both raw and preprocessed data from the HCP-YA dataset are available through the official website (https://www.humanconnectome.org/study/hcp-young-adult/document/1200-subjects-data-release). Human gene expression data are available from the Allen Brain Atlas (https://human.brainmap.org) and the receptor density atlas is available at https://github.com/netneurolab/hansen_receptors. The PLS1 genes and enriched terms of the cell-type-specific genes are provided in Supplementary Data 1. All data supporting the findings of this study are included in the paper and its Supplementary Information, and all additional information is available from the authors upon reasonable request.

## Code availability

All the codes used to generate the results will be available on GitHub (https://github.com/TianyiYanLab/CUD_Pathology_Modelling).

## Acknowledgements

The authors thank the SUDMEX team for their support and help. This work was supported by the National Natural Science Foundation of China (grant numbers 62336002, 62406025 and 82302175); the STI 2030-Major Projects (grant number 2022ZD0208500); the National Science and Technology Innovation 2030 Program (grant number 2021ZD0200500); the Beijing Nova Program (grant number 20230484465); and the BIT Research and Innovation Promoting Project (grant number 2024YCXZ027).

