## SupplementaryInformation for "Structural connectome architecture and biological vulnerability shape cortical atrophy in cocaine use disorder"

Han *et al.*

### Supplementary Figure 1

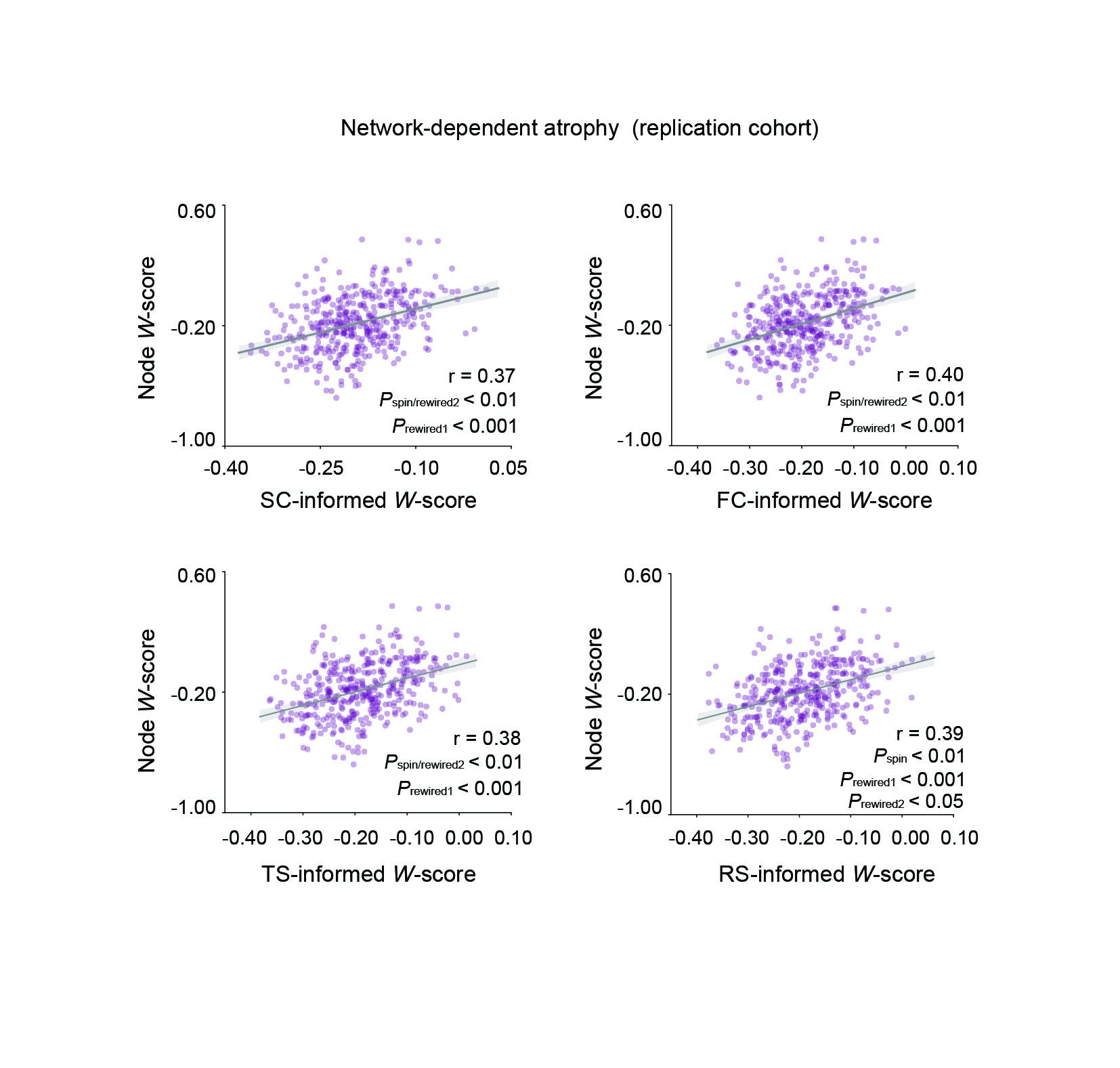

**Fig. S1** **Network-dependent cortical atrophy in the replication cohort.** The region-to-neighbour atrophy correlations were significant across the structural connectome, functional connectome, transcriptomic and receptor similarity networks in the replication cohort. The significance of correlations was evaluated against the spin and rewired null models (1, 000 iterations). SC, structural connectome; FC, functional connectome; TS, transcriptomic similarity; RS, receptor similarity.

### Supplementary Figure 2

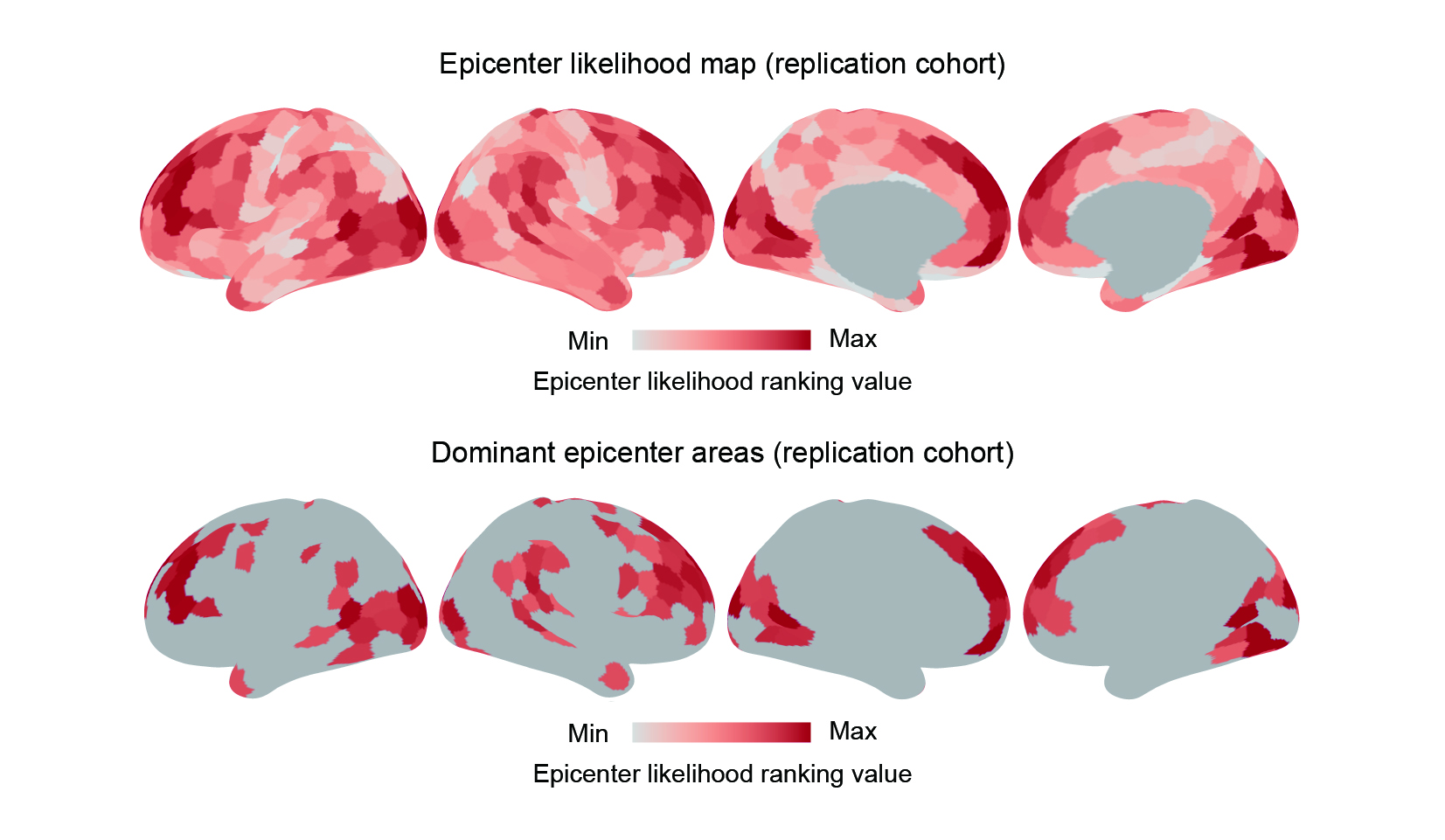

**Fig. S2** **Epicenter identification in the replication cohort.** Regions were ranked based on their atrophy values and disease exposure values, and the average of the two ranks was used to represent the probability of a region being an epicenter. The regions with the top 25% of epicenter likelihood rankings were depicted in the bottom panel.

### Supplementary Figure 3

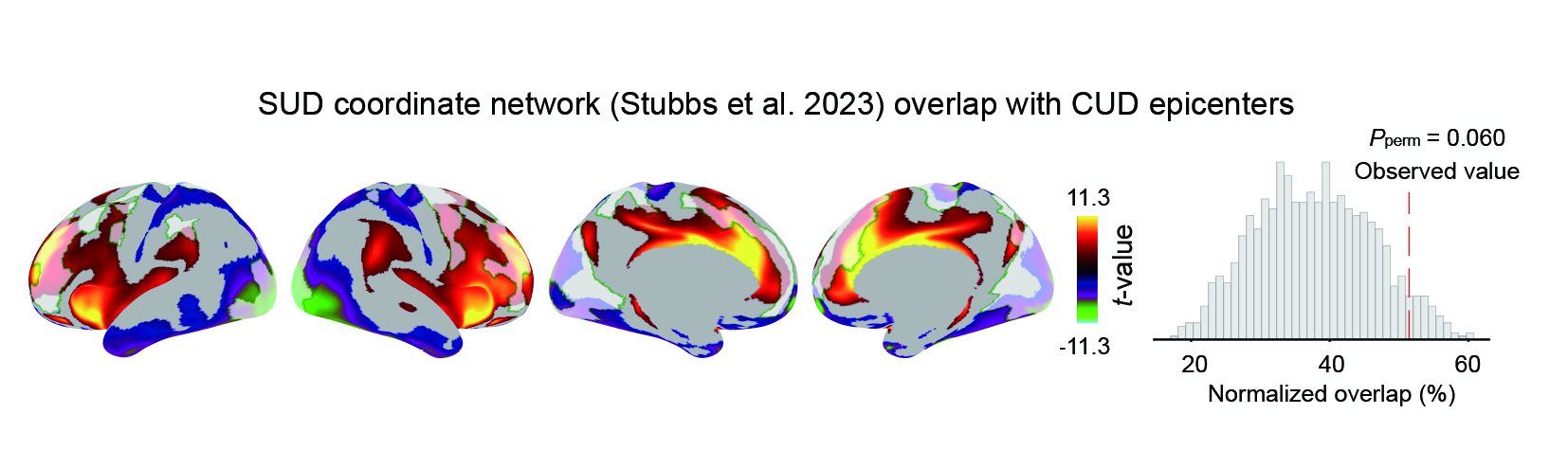

**Fig. S3** **Epicenter validation.** The top 25% of regions with the highest epicenter likelihood in CUD were part of the SUD network and showed greater overlap (*P_perm_* = 0.060) with this network compared to a null distribution generated by randomly rotating these top epicenter regions on the cortical surface (spin tests; 1,000 iterations). SUD, substance use disorder.

### Supplementary Figure 4

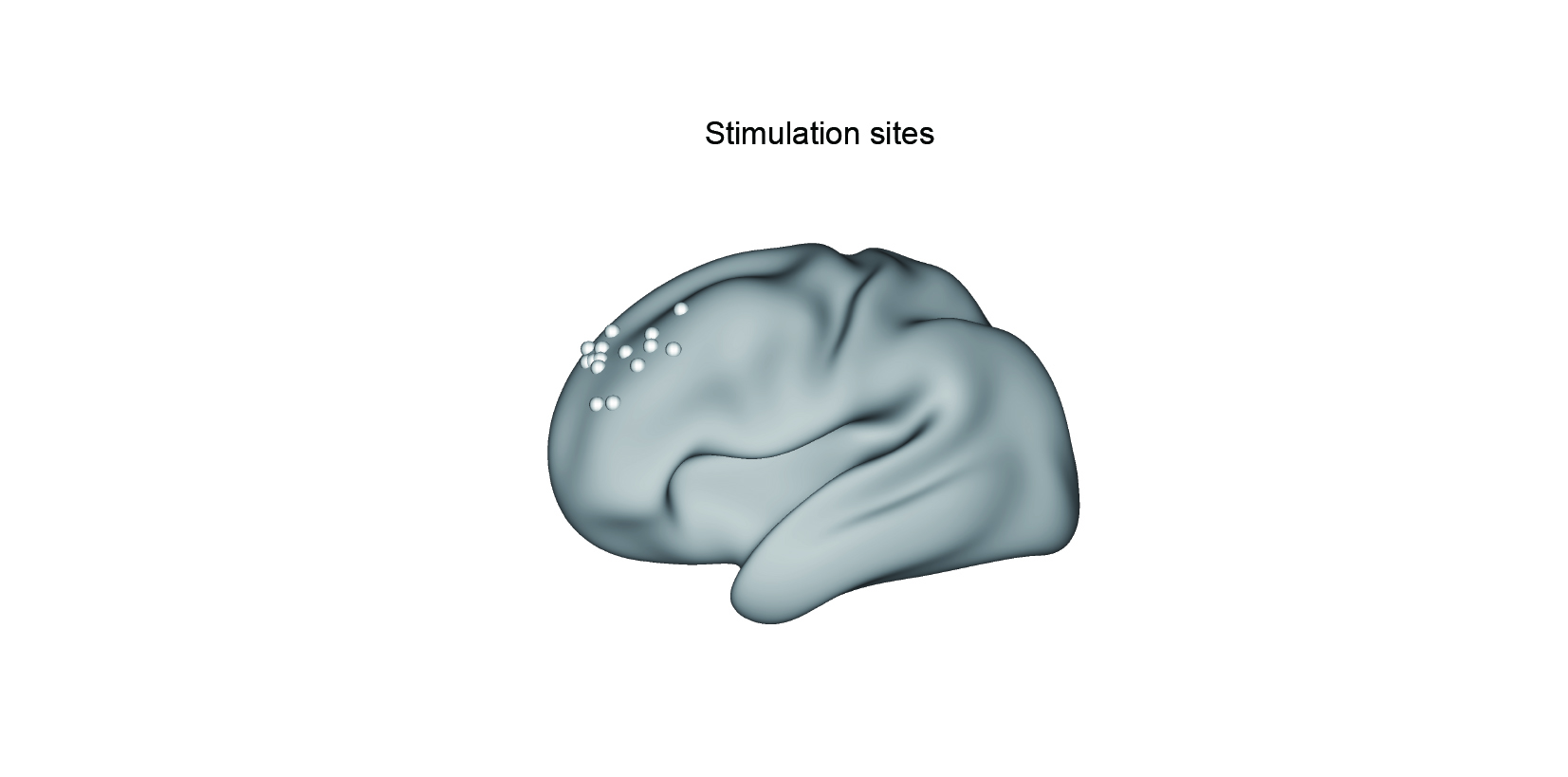

**Fig. S4 Spatial distribution of rTMS sites in the discovery cohort.** All patients (N=15, age range: 25-47 years, 2 females) from the active group of SUDMEX-TMS cohort with both stimulation target localization information and functional magnetic resonance imaging (fMRI) data were included to generate the rTMS response map. rTMS, repeated transcranial magnetic stimulation.

### Supplementary Figure 5

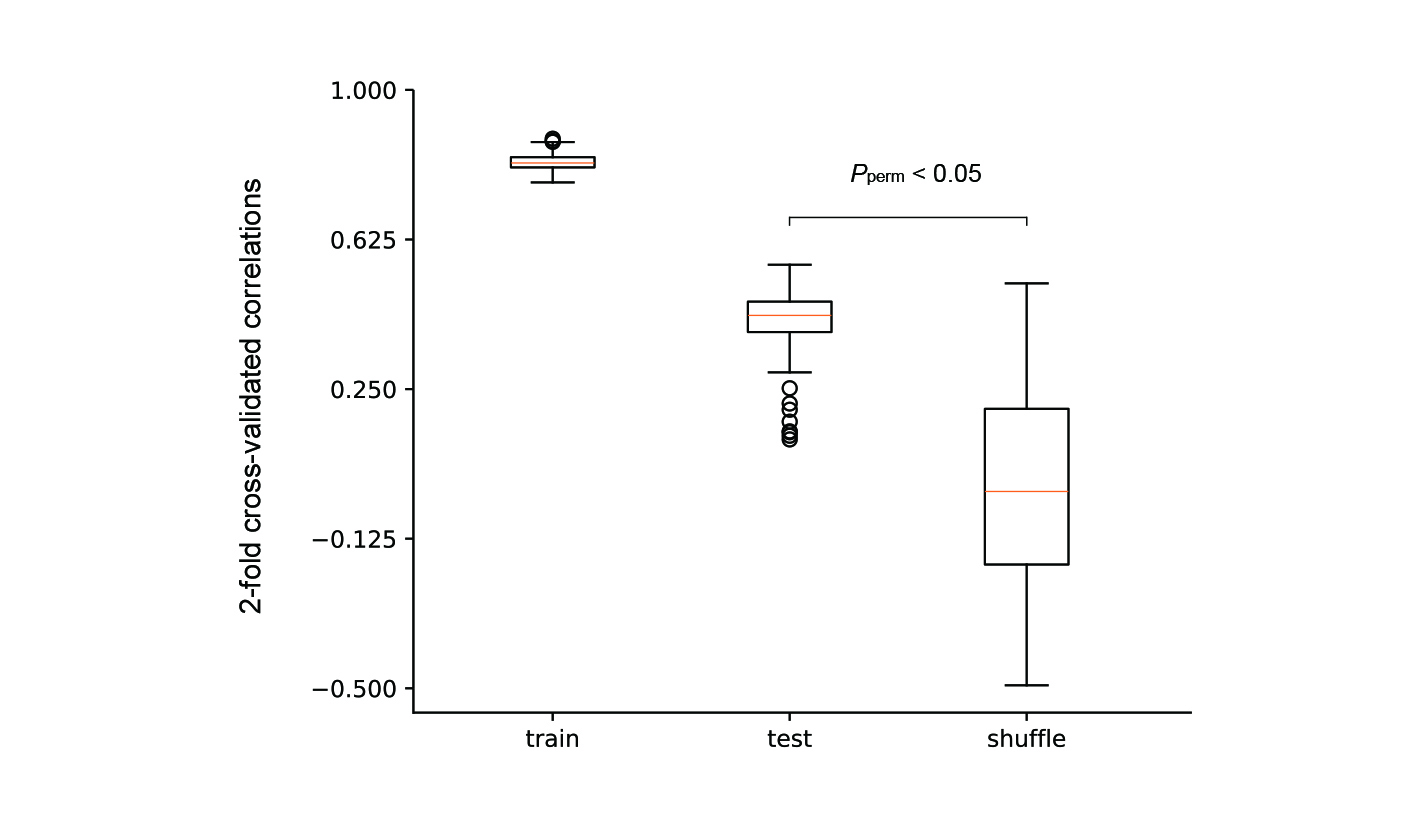

**Fig. S5** **Cross-validation of the PLS analysis.** 2-fold cross-validation was used to evaluate the out-of-sample multivariate correlation between cortical epicenter patterns and individual behavioural features. In each iteration, the dataset was randomly split into equal-sized training and testing subsets. Partial least squares (PLS) regression was applied to the training set to derive singular vector weights, which were then used to project the test data and compute patient-specific scores. The correlation between predicted and actual behavioural features in the test set was calculated, and this process was repeated 150 times to generate a distribution of out-of-sample correlation values. To assess the statistical significance of this distribution, we additionally performed 150 permutation tests by shuffling the rows of the epicenter matrix and repeating the cross-validation procedure, thereby generating a null distribution of correlation values. A non-parametric *P* value (*P* = 0.027) was obtained by comparing the median of empirical correlations to the null distribution.

### Supplementary Figure 6

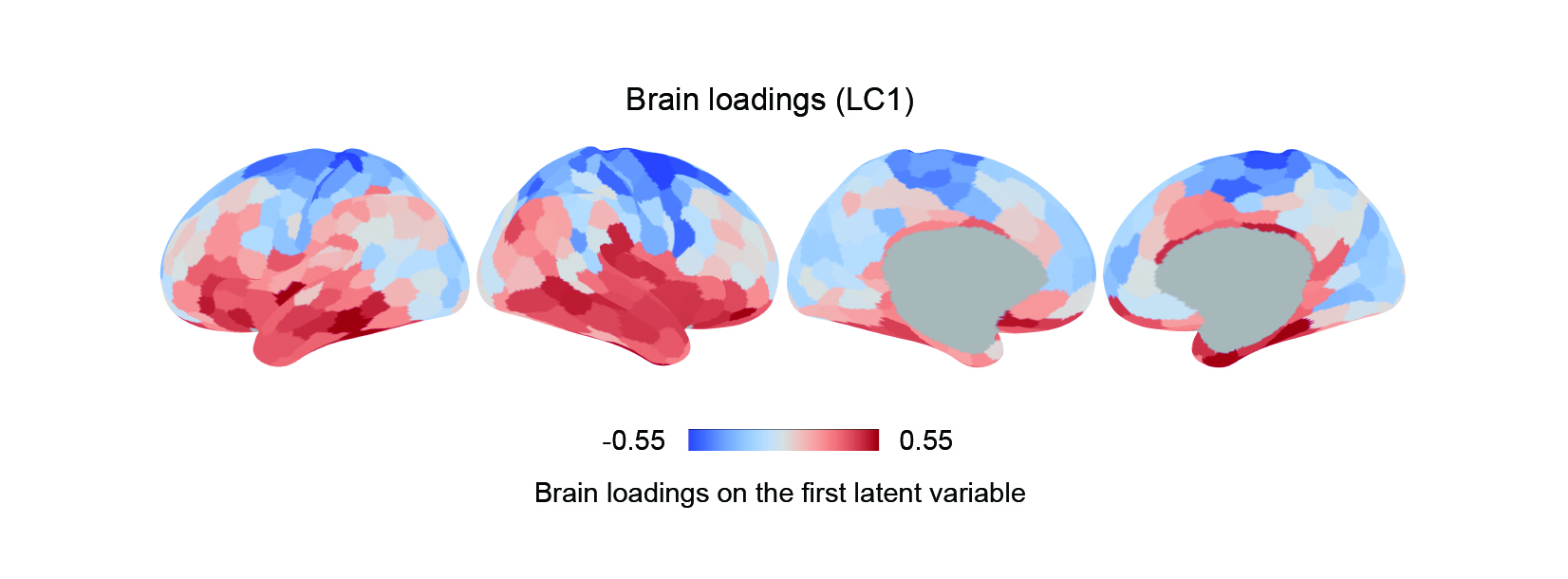

**Fig. S6 Relating cortical epicenter patterns with individual behavioural features.** Brain loadings of the first latent component (LC1) from a PLS analysis. Red (or blue) colour indicates that greater epicenter probability is positively (or negatively) associated with LC1.

### Supplementary Figure 7

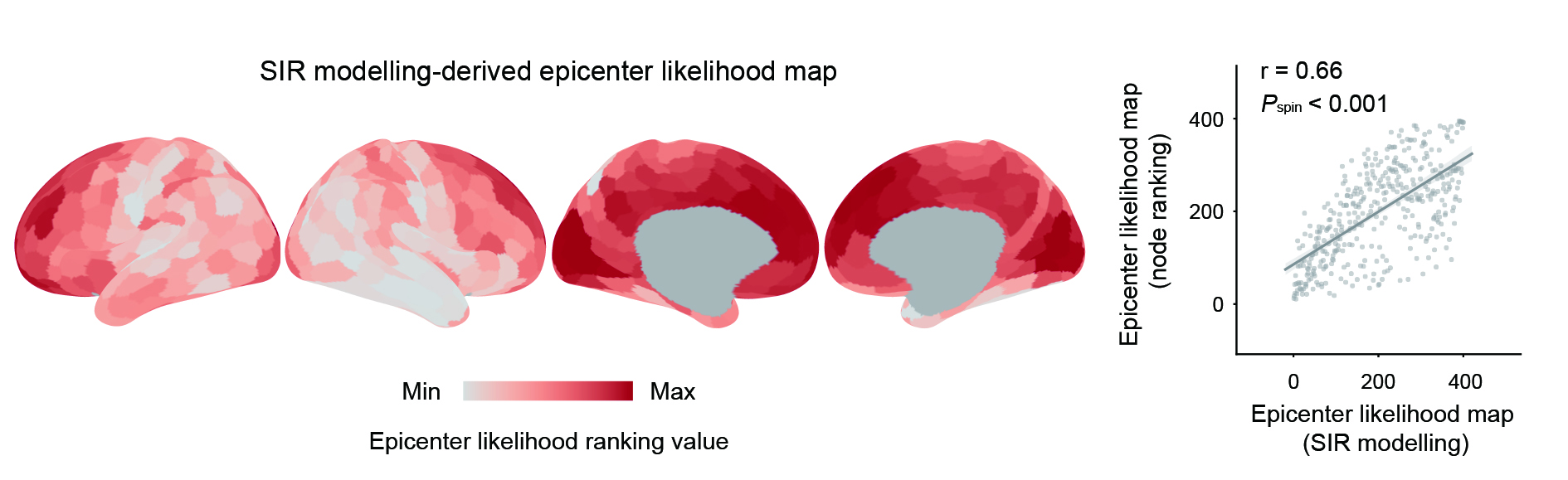

**Fig. S7 Epicenter identification using an agent-based Susceptible-Infected-Removed (SIR) model in the discovery cohort.** In brief, the model simulated the propagation of misfolded pathogenic proteins through the brain network. Epicenter likelihood maps derived from the data-driven ranking-based method and SIR method exhibited significantly correlated spatial distribution patterns (r = 0.66, *P*_spin_ < 0.001).

### Supplementary Figure 8

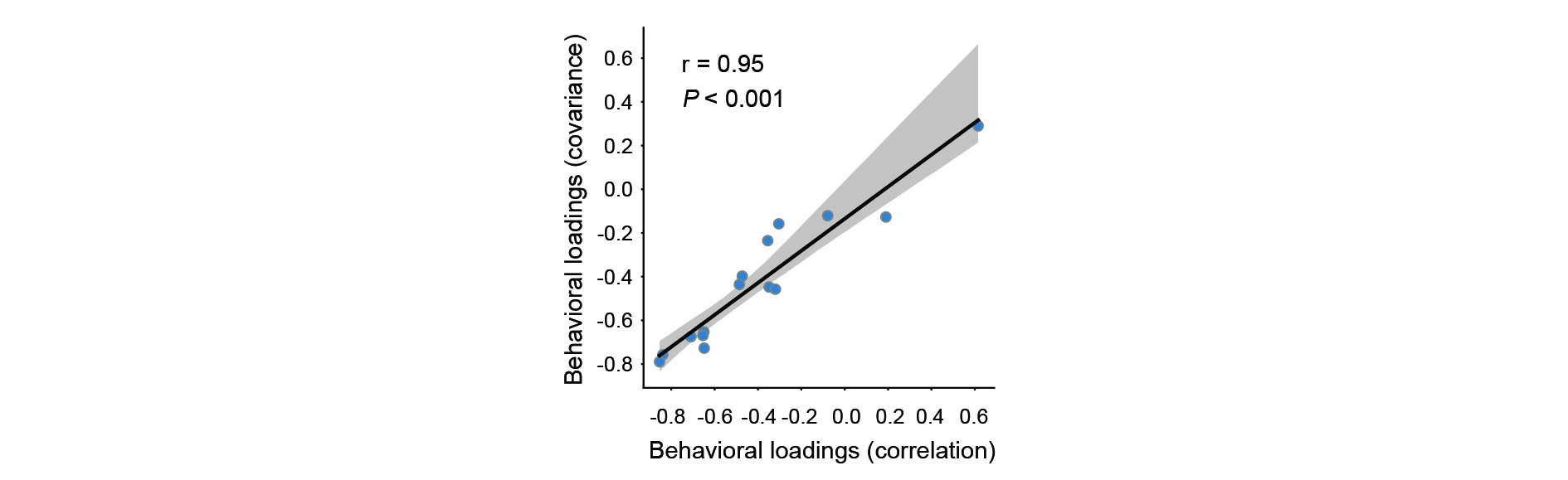

**Fig. S8 Consistency of behavioural loadings derived from covariance and correlation-based analyses.** Scatter plot demonstrating the high consistency (r = 0.95, *P* < 0.001) between behavioural loadings on the first latent component (LC1) obtained using covariance and correlation-based analytical approaches. Each dot represents a behavioural measure.
